## Supplementary material for "iPSC-derived models of PACS1 syndrome reveal transcriptional and functional deficits in neuron activity": Fig. S

**This file includes:**

Figures S1-S15 and legends

Table S1 and legend

Table S2 legend (separate file)

#### SUPPLEMENTARY FIGURES

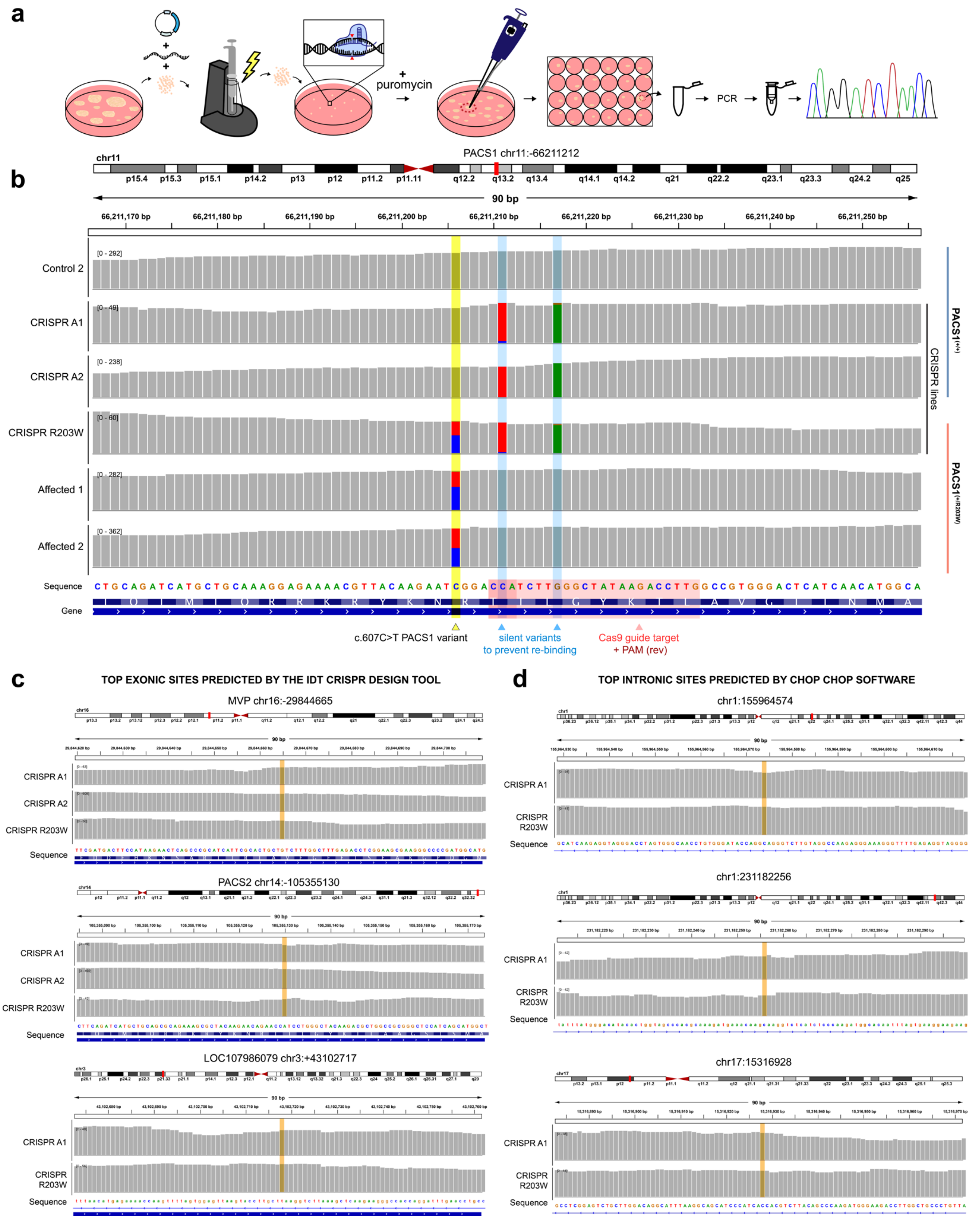

**Figure S1. Generation and validation of isogenic stem cell lines. a)** iPSCs were electroporated with a plasmid expressing Cas9, a guide targeting exon 4 of *PACS1*, and an ssODN containing the c.607C>T variant in addition to two silent mutations (c.612C>T and c.618G>A) to prevent re-binding of the guide. **b**) Sequencing validates introduction of the heterozygous c.607C>T variant in the CRISPR R203W line - also observed in patient lines A1 and A2 (boxed in yellow) - and successful correction of the patient variant in CRISPR A1 and CRISPR A2 lines. In CRISPR-edited lines, the two downstream additional variants (boxed in blue) are the silent mutations. The guide target region is boxed in red with the PAM site towards the 5' end. **c)** Top predicted exonic off-target sites predicted by the IDT CRISPR design tool. The predicted site is highlighted in orange with the surrounding 90 bp showing no unintended edits were introduced. CRISPR A2 is missing from the final panel as it received exome sequencing instead of whole-genome sequencing, which did not cover this noncoding RNA. **d)** Top predicted intronic off-target sites predicted by CHOP CHOP software. Not all lines are present as only CRISPR R203W and CRISPR A1 received whole-genome sequencing.

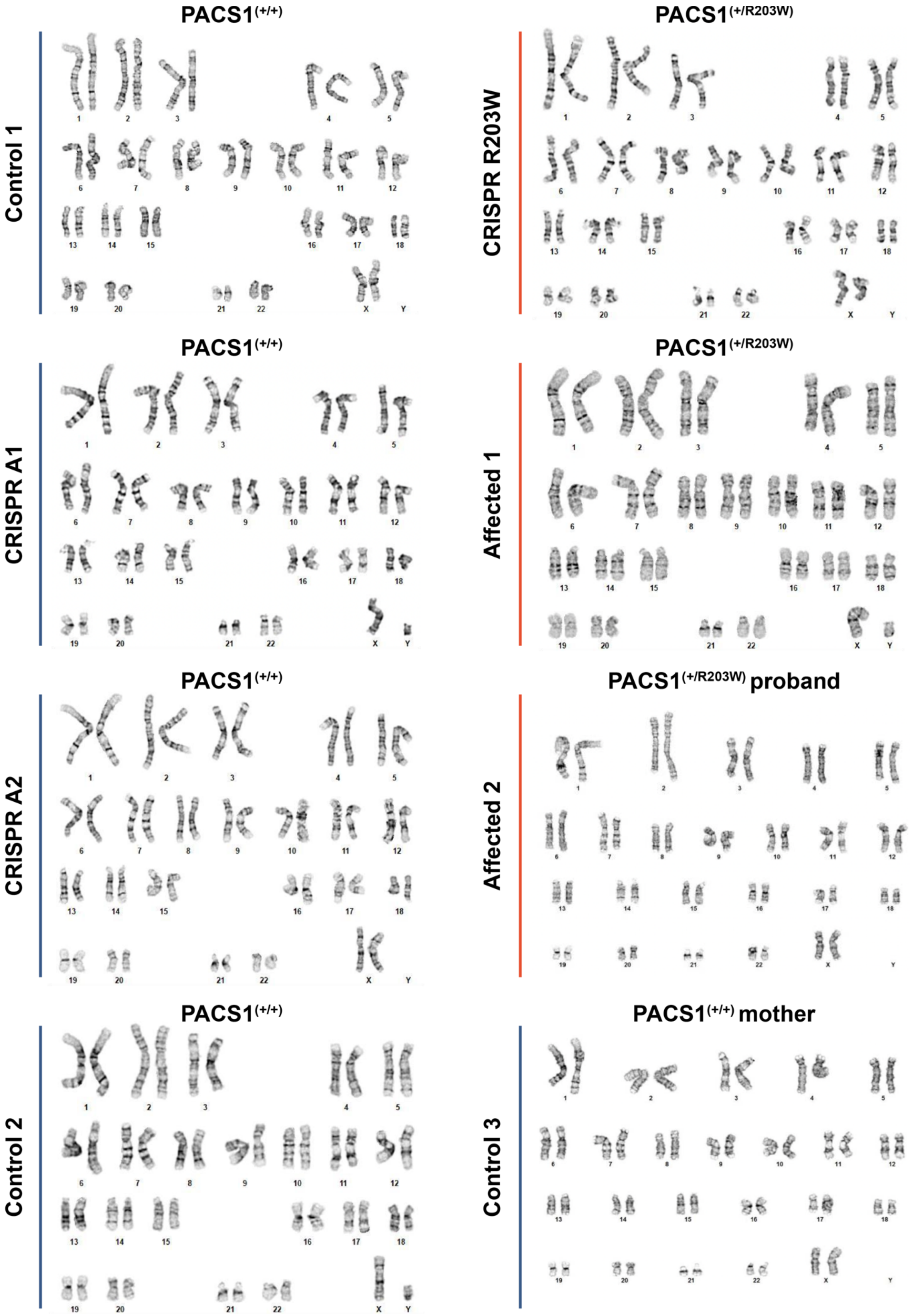


**Figure S2. iPSC parent and CRISPR-edited lines display normal karyotypes.** Karyotype analysis for all iPSC lines used in this study. PACS1$^{(+/+)}$ and PACS1$^{(+/R203W)}$ lines display a normal karyotype, including CRISPR-edited lines.

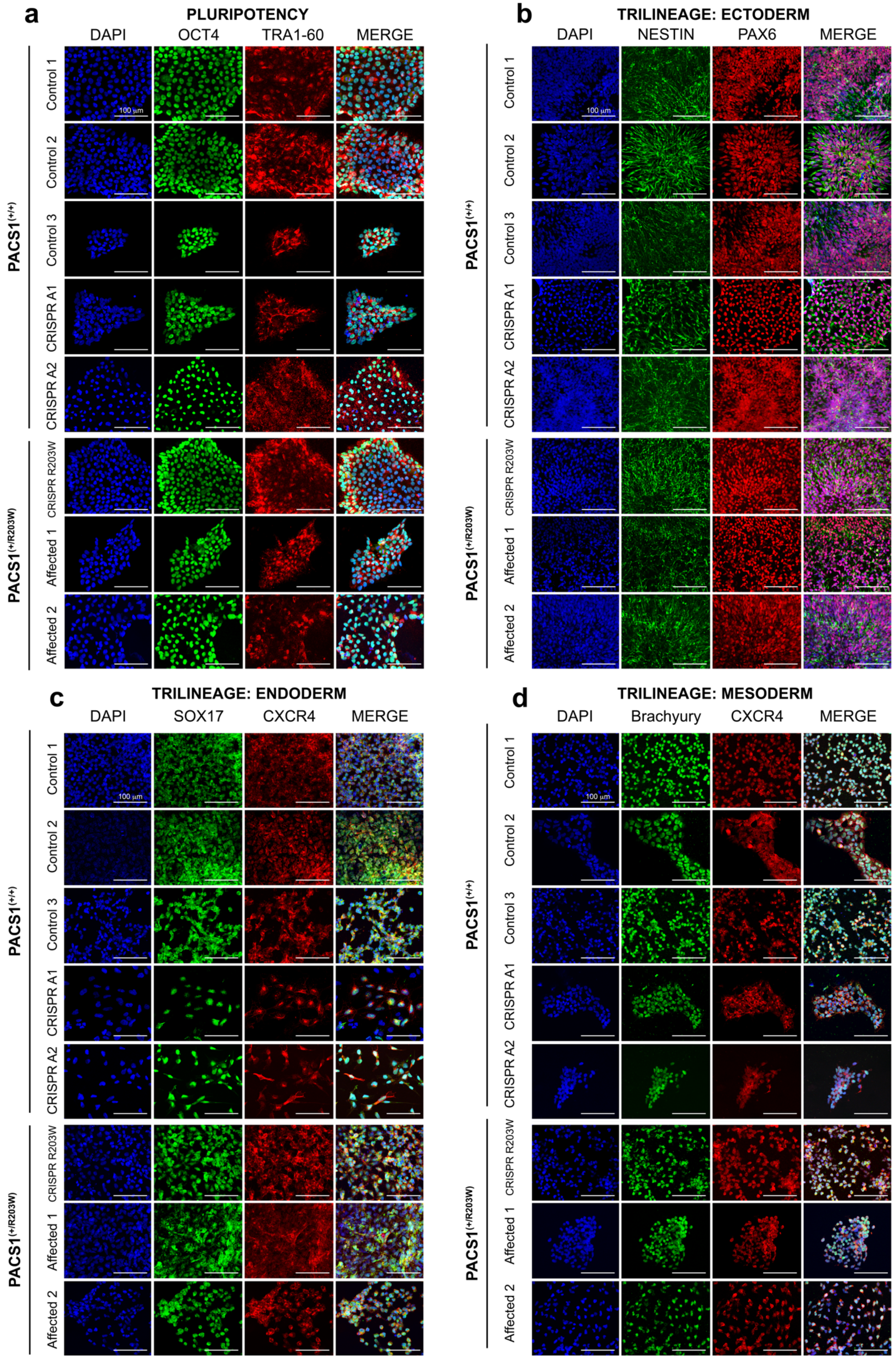
a
PLURIPOTENCY
DAPI
OCT4
TRA1-60
MERGE
b
TRILINEAGE: ECTODERM
DAPI
NESTIN
PAX6
MERGE
c
TRILINEAGE: ENDODERM
DAPI
SOX17
CXCR4
MERGE
d
TRILINEAGE: MESODERM
DAPI
Brachyury
CXCR4
MERGE
PACS1(+/+)
PACS1(+/R203W)
Control 1
Control 2
Control 3
CRISPR A1
CRISPR A2
CRISPR R203W
Affected 1
Affected 2
100 µm

**Figure S3: iPSC parent and CRISPR-edited lines express pluripotent markers and have the capacity to generate all three germ layers. a)** Immunohistochemistry analysis demonstrates iPSC lines express pluripotent markers OCT4 and TRA1-60. Scale bar = 100 µm. Pluripotent capacity is further demonstrated by the generation of all three germ layers in **b-d. b)** iPSCs differentiated to ectodermal lineage as indicated by co-expression of PAX6 and Nestin. Scale bar = 100 µm. **c)** iPSCs directed to endodermal lineage as indicated by co-expression of SOX17 and CXCR4. Scale bar = 100 µm. **d)** iPSCs differentiated towards mesodermal lineage as indicated by co-expression of Brachyury and CXCR4. Scale bar = 100 µm.

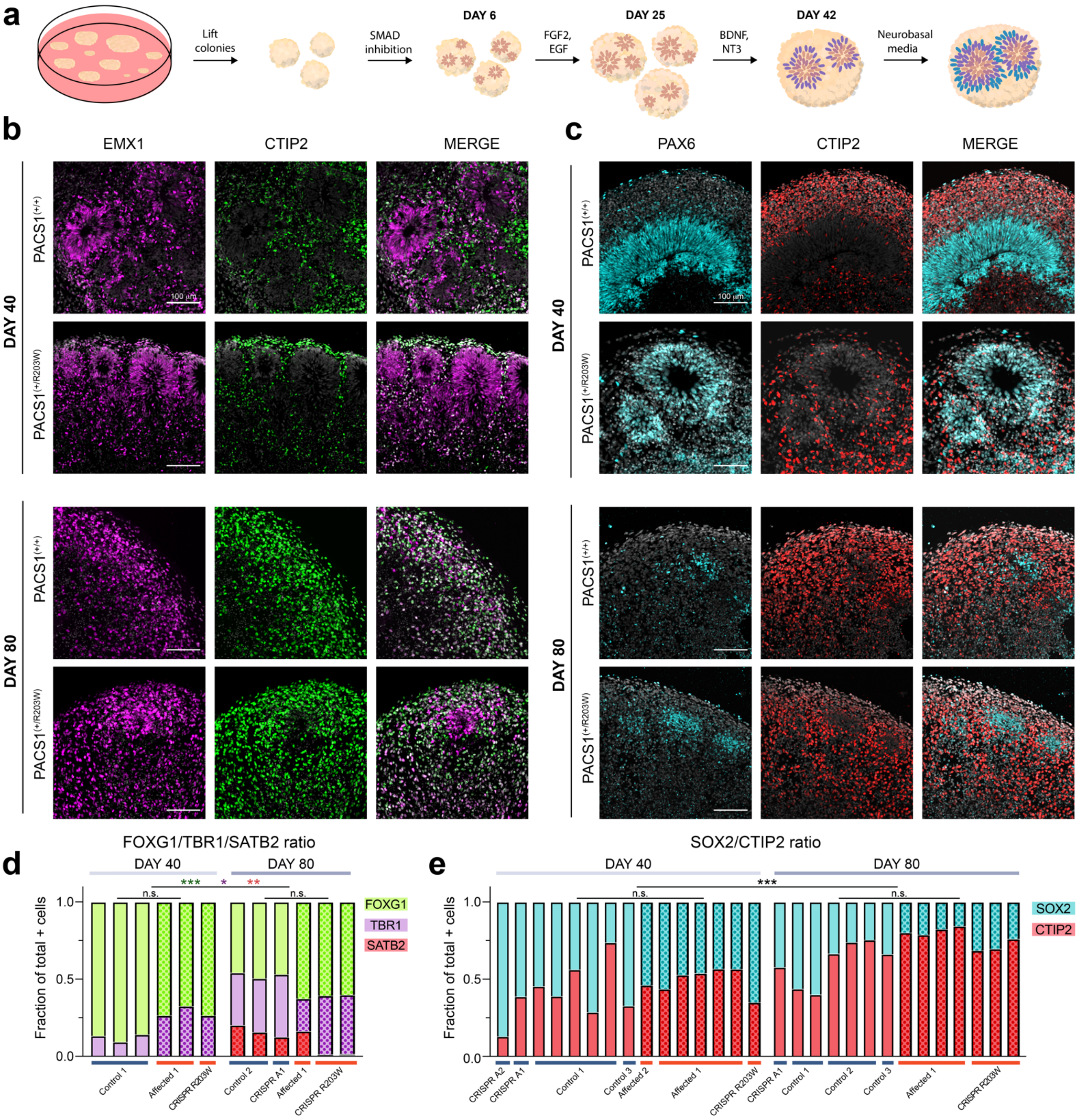

**Figure S4. Generation and validation of a dorsal cortical organoid model system**. **a)** Dorsal cortical organoids are generated using a growth factor program to induce neuroectoderm differentiation, proliferation, and survival. **b)** Expression patterns of dorsal forebrain markers EMX1 and CTIP2 in day 40 and 80 organoids. Scale bar = 100 µm. DAPI is in greyscale. **c)** Expression patterns of dorsal forebrain progenitor protein PAX6 and postmitotic neuron marker CTIP2 in day 40 and 80 organoids. Scale bar = 100 µm. DAPI is in greyscale. **d**) FOXG1/TBR1/SATB2 ratios shown in **Fig. 1d** by organoid. Bar shows the mean fraction of positive cells contributing to the total number of positive cells per image field. Stains are different by day (FOXG1 $F_{1,9}$=24.004, $p_{adj}$ = 8.43E-04; TBR1 $F_{1,9}$=9.459, $p_{adj}$ = 0.013; SATB2 $H_1$=9.466, $p_{adj}$ = 0.004) but not genotype (FOXG1 $F_{1,9}$=0.061, $p_{adj}$ =0.811; TBR1 $F_{1,9}$=1.811, $p_{adj}$ = 0.211; SATB2 $H_1$ = 0.183, $p_{adj}$=0.669) using a two-way ANOVA unless $p < 0.05$ in a Shapiro test of normality (see Methods). $n$ = 3 organoids for each condition (Day 40: three C1, one CRISPR R203W, and two A1 organoids from four independent differentiations; Day 80: two C2, one CRISPR A1, two CRISPR R203W, and one A1 organoids from three independent differentiations. See **Table S1**). Data from PACS1$^{(+/R203W)}$ lines is indicated by the dot pattern. **e)** SOX2/CTIP2 ratios shown in **Fig. 1c** by organoid. Bar shows the mean fraction of positive cells contributing to the total number of positive cells per image field. Mean ± SEM fraction of CTIP2+ or SOX2+ cells contributing to the total number of positive cells per field of view. The ratio was significant between time points ($H_1$=12.808, $p_{adj}$ = 6.90E-04) but not genotypes ($H_1$=3.360, $p_{adj}$ = 0.067). $n$ = 7+ organoids per condition (Day 40: five C1, one C3, one CRISPR A1, one CRISPR A2, one CRISPR R203W, five A1, and one A2 organoids from four independent differentiations; Day 80: two C1, three C2, one C3, one CRISPR A1, three CRISPR R203W, and four A1 organoids from three independent differentiations. See **Table S1**). Data from PACS1$^{(+/R203W)}$ lines is indicated by the dot pattern.

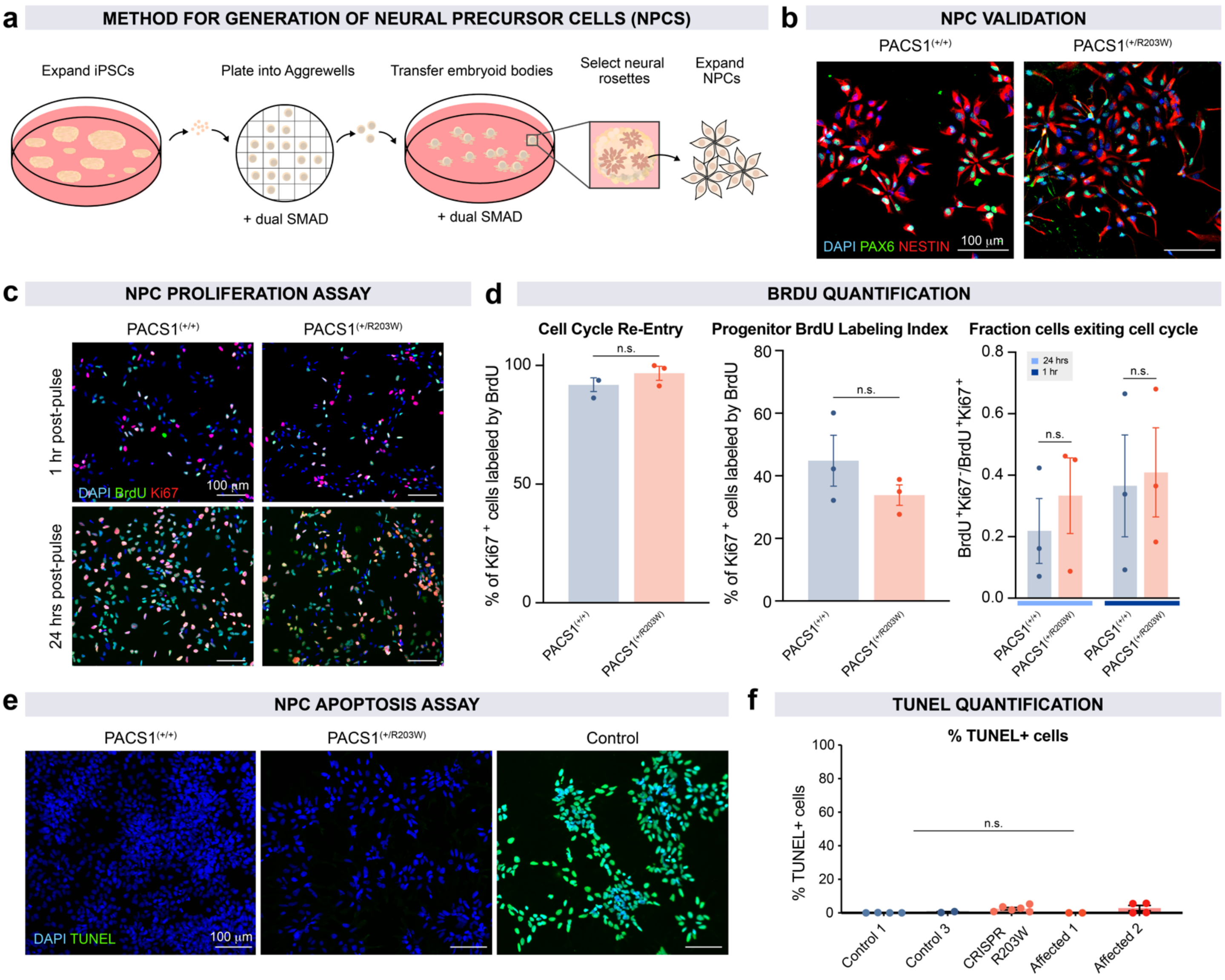


**Figure S5: PACS1 p.R203W variant does not impact neural precursor cell (NPC) proliferation or apoptosis rates. a)** Protocol for the differentiation of iPSCs to NPCs. In brief, iPSCs are dissociated and plated in Aggrewell plates to generate embryoid bodies (EBs). EBs are exposed to dual SMAD inhibitors, encouraging ectoderm formation. Rosettes are selected and expanded to generate a pure NPC population. **b)** High-quality NPCs were generated in both genotypes, as indicated by co-expression of ectoderm markers PAX6 and Nestin. Scale bar = 100 µm. **c)** Proliferation rate of NPCs was assessed by pulsing the thymidine analog BrdU and co-staining for mitotic marker Ki67. Scale bar = 100 µm. Resulting metrics are calculated in **d)**. Mean ± SEM of proliferation metrics as calculated from the BrdU and Ki67 immunohistochemistry images in **c**. *n* = 3 biological replicates from two independent experiments (one Cl, one C2, one C3, one CRISPR R203W, one A1, and one A2 coverslip average). Cell cycle re-entry was indicated by the percentage of Ki67+ cells labeled with BrdU after 24 hours. Genotype had no effect on cell cycle re-entry ($t_{4.00}$ = -1.282, $p$ = 0.269 calculated using a Welch Two Sample two-sided t test). The progenitor BrdU labeling index is calculated by the percentage of Ki67+ progenitors that also incorporated BrdU and is not different between genotypes ($t_{2.62}$=1.253, $p$ = 0.310 using a Welch Two Sample two-sided t test). Cells exiting the cell cycle are indicated by number of BrdU+ cells minus double-positive cells over total double-positive cells and is not different between genotypes ($F_1$=0.376, $p_{adj}$ = 0.555 using a two-way ANOVA with a Tukey HSD correction) . **e)** NPC apoptosis rate was assessed using a TUNEL assay. Co-localization of DAPI and TUNEL is shown in PACS1$^{(+/+)}$ and PACS1$^{(+/R203W)}$ NPCs, along with a control in which PACS1$^{(+/R203W)}$ cells were treated with DNAse I for 30 minutes. Scale bar = 100 µm. **f)** Quantification of the TUNEL assay in **e** shows a very low rate of TUNEL+ cells in each cell line tested. Genotype has no effect on the incidence of TUNEL+ cells ($t_{2.03}$ = 1.803, $p$ = 0.211 calculated using a Welch Two Sample two-sided t test).

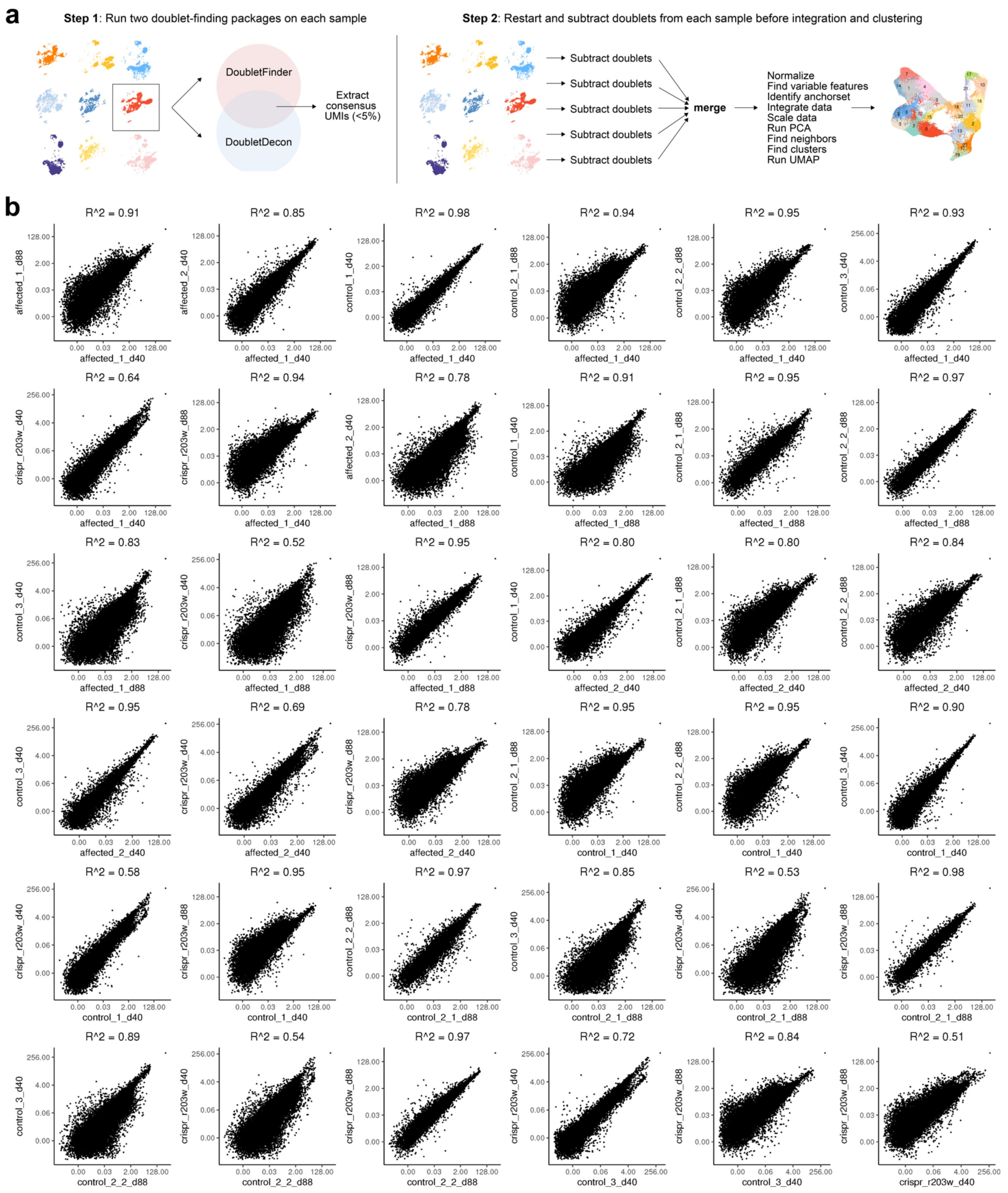


**Figure S6. Single-cell RNA sequencing in day 40 and 88 organoids. a)** Initial sample processing schematic. First, each sample was individually constructed, and two separate doublet-finding packages were utilized.[83,]

[84] Consensus doublet barcodes were removed, and the samples were then all processed in parallel. **b)** Pearson's correlation of gene expression between all sample pair combinations from nine organoids total across four independent differentiations (Day 40: one C1, one C3, one CRISPR R203W, one A1, and one A2 organoid; Day 88: two C2, one CRISPR R203W, and one A1 organoid. See **Table S1**).

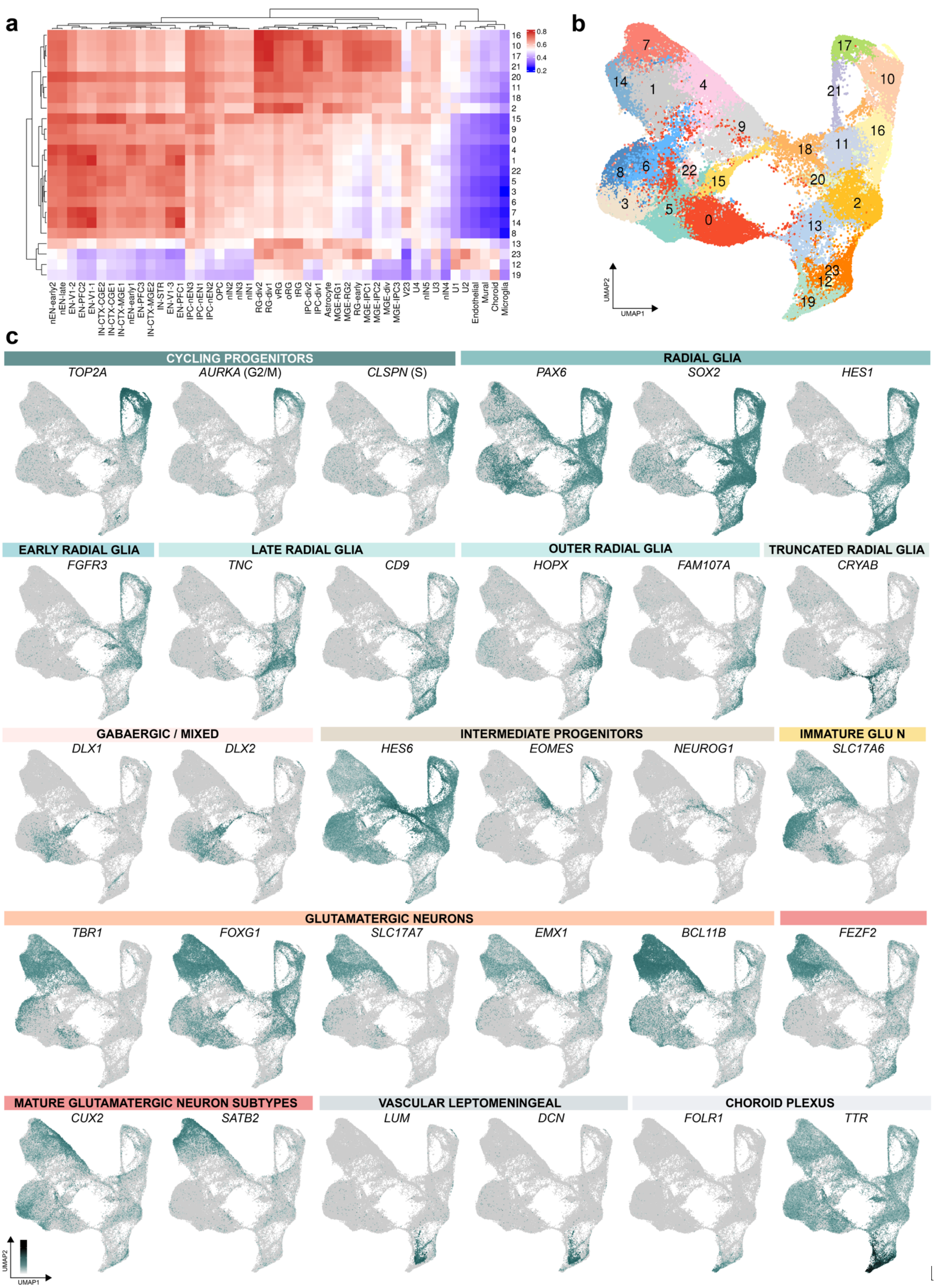
a
16
10
17
21
20
11
18
2
15
9
0
4
1
22
5
3
6
7
14
8
13
23
12
19
0.8
0.6
0.4
0.2
nEN-early2
nEN-late
EN-V1-2
EN-PFC2
EN-V1-1
IN-CTX-CGE2
IN-CTX-CGE1
IN-CTX-MGE1
nEN-early1
EN-PFC3
IN-CTX-MGE2
IN-STR
EN-V1-3
EN-PFC1
IPC-nEN3
IPC-nEN1
IPC-nEN2
OPC
nIN2
nIN3
nIN1
RG-div2
RG-div1
vRG
oRG
tRG
IPC-div2
IPC-div1
Astrocyte
MGE-RG1
MGE-RG2
RG-early
MGE-IPC1
MGE-IPC2
MGE-div
MGE-IPC3
V23
U4
nIN5
U3
nIN4
U1
U2
Endothelial
Mural
Choroid
Microglia
b
UMAP2
UMAP1
c
CYCLING PROGENITORS
TOP2A
AURKA (G2/M)
CLSPN (S)
RADIAL GLIA
PAX6
SOX2
HES1
EARLY RADIAL GLIA
FGFR3
LATE RADIAL GLIA
TNC
CD9
OUTER RADIAL GLIA
HOPX
FAM107A
TRUNCATED RADIAL GLIA
CRYAB
GABAERGIC / MIXED
DLX1
DLX2
INTERMEDIATE PROGENITORS
HES6
EOMES
NEUROG1
IMMATURE GLU N
SLC17A6
GLUTAMATERGIC NEURONS
TBR1
FOXG1
SLC17A7
EMX1
BCL11B
FEZF2
MATURE GLUTAMATERGIC NEURON SUBTYPES
CUX2
SATB2
VASCULAR LEPTOMENINGEAL
LUM
DCN
CHOROID PLEXUS
FOLR1
TTR
UMAP2
UMAP1

**Figure S7. Classification of cell types in single-cell RNA sequencing data. a)** Cell type annotation was bioinformatically aided with the Clustifyr[85] package by correlation of the transcriptome of each Seurat cluster to a reference fetal tissue dataset published by Nowakowski *et al.*[86] **b)** Uniform manifold approximation and projection (UMAP) spatial location of the corresponding clusters in **a**. Data in **b-c** is shown from nine organoids total across four independent differentiations (Day 40: one C1, one C3, one CRISPR R203W, one A1, and one A2 organoid; Day 88: two C2, one CRISPR R203W, and one A1 organoid. See **Table S1**). **c)** Feature expression for all cells in UMAP space of additional canonical cell type markers. *TOP2A* shows actively dividing cells with *AURKA* specific to G2/M phase and *CLSPN* to S phase; *PAX6, SOX2,* and *HES1* denote radial glia populations; *FGFR3* marks early radial glia groups; *TNC* and *CD9* are enriched in late radial glia; *HOPX* and *FAM107A* are enriched in outer radial glia; *CRYAB* is strongly expressed in the truncated radial glia group; *DLX1* and *DLX2* expression are highest in the inhibitory neurons, but also lowly expressed in the mixed immature neuron population; IPCs are determined by the colocalization of *HES6, EOMES,* and *NEUROG1*; immature glutamatergic neurons express *SLC17A6*; and glutamatergic neurons express *TBR1, FOXG1, SLC17A7, EMX1,* and *BCL11B*. Mature glutamatergic neuron groups are further defined by subgroups with high expression of subcortical projection marker *FEZF2,* cortical layer II-III marker *CUX2*; or upper layer marker *SATB2.* Small subpopulations express late radial glia transcripts, but more highly express markers of vascular leptomeningeal cells such as *LUM* and *DCN*; or choroid plexus markers such as *FOLR1* and *TTR.* Scale is in units of $\log_{10}$(value + 0.1) and extends from 0-2 for all features except *HES6* and *TTR*, which extend from 0-3.

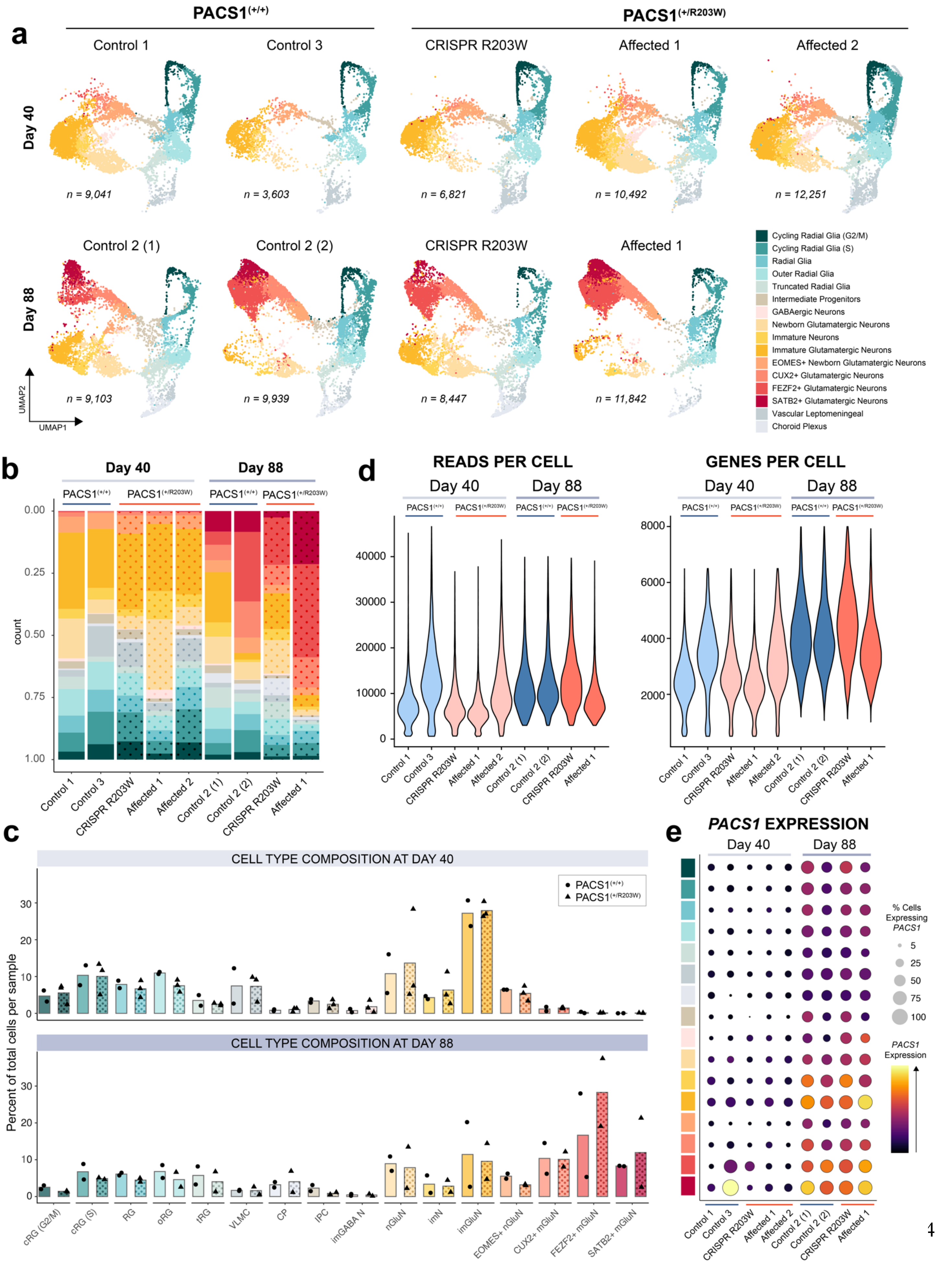

a
PACS1(+/+)
PACS1(+/R203W)
Control 1
Control 3
CRISPR R203W
Affected 1
Affected 2
Day 40
n = 9,041
n = 3,603
n = 6,821
n = 10,492
n = 12,251
Control 2 (1)
Control 2 (2)
CRISPR R203W
Affected 1
Day 88
n = 9,103
n = 9,939
n = 8,447
n = 11,842
UMAP2
UMAP1
Cycling Radial Glia (G2/M)
Cycling Radial Glia (S)
Radial Glia
Outer Radial Glia
Truncated Radial Glia
Intermediate Progenitors
GABAergic Neurons
Newborn Glutamatergic Neurons
Immature Neurons
Immature Glutamatergic Neurons
EOMES+ Newborn Glutamatergic Neurons
CUX2+ Glutamatergic Neurons
FEZF2+ Glutamatergic Neurons
SATB2+ Glutamatergic Neurons
Vascular Leptomeningeal
Choroid Plexus
b
Day 40
Day 88
PACS1(+/+)
PACS1(+/R203W)
count
0.00
0.25
0.50
0.75
1.00
Control 1
Control 3
CRISPR R203W
Affected 1
Affected 2
Control 2 (1)
Control 2 (2)
CRISPR R203W
Affected 1
d
READS PER CELL
GENES PER CELL
Day 40
Day 88
0
10000
20000
30000
40000
2000
4000
6000
8000
c
CELL TYPE COMPOSITION AT DAY 40
CELL TYPE COMPOSITION AT DAY 88
PACS1(+/+)
PACS1(+/R203W)
Percent of total cells per sample
0
10
20
30
cRG (G2/M)
cRG (S)
RG
oRG
tRG
VLMC
CP
IPC
imGABA N
nGluN
imN
imGluN
EOMES+ nGluN
CUX2+ mGluN
FEZF2+ mGluN
SATB2+ mGluN
e
PACS1 EXPRESSION
% Cells Expressing PACS1
5
25
50
75
100
PACS1 Expression

**Figure S8. Distribution of cell types across day 40 and day 88 single-cell RNA sequencing samples. a)** UMAP plot of each sample colored according to cell type. Data in **a-e** is shown from nine organoids total across four independent differentiations (Day 40: one C1, one C3, one CRISPR R203W, one A1, and one A2 organoid; Day 88: two C2, one CRISPR R203W, and one A1 organoid. See Table S1). **b)** Bar chart of the distribution of cell type proportions between all nine samples. Data from PACS1$^{(+/R203W)}$ lines is indicated by the dot pattern. **c)** Quantification of the bar chart in **b** showing the percentage contribution of each cell type to PACS1$^{(+/+)}$ and PACS1$^{(+/R203W)}$ samples at each time point. Data from PACS1$^{(+/R203W)}$ lines is indicated by the dot pattern. $p > 0.05$ for each condition. **d)** Violin plot of reads and genes per cell for each sample. **e)** Dot plot of *PACS1* expression for all cell types and samples. Yellow color indicates highest normalized expression and size indicates the percentage of cells in the group that express *PACS1*.

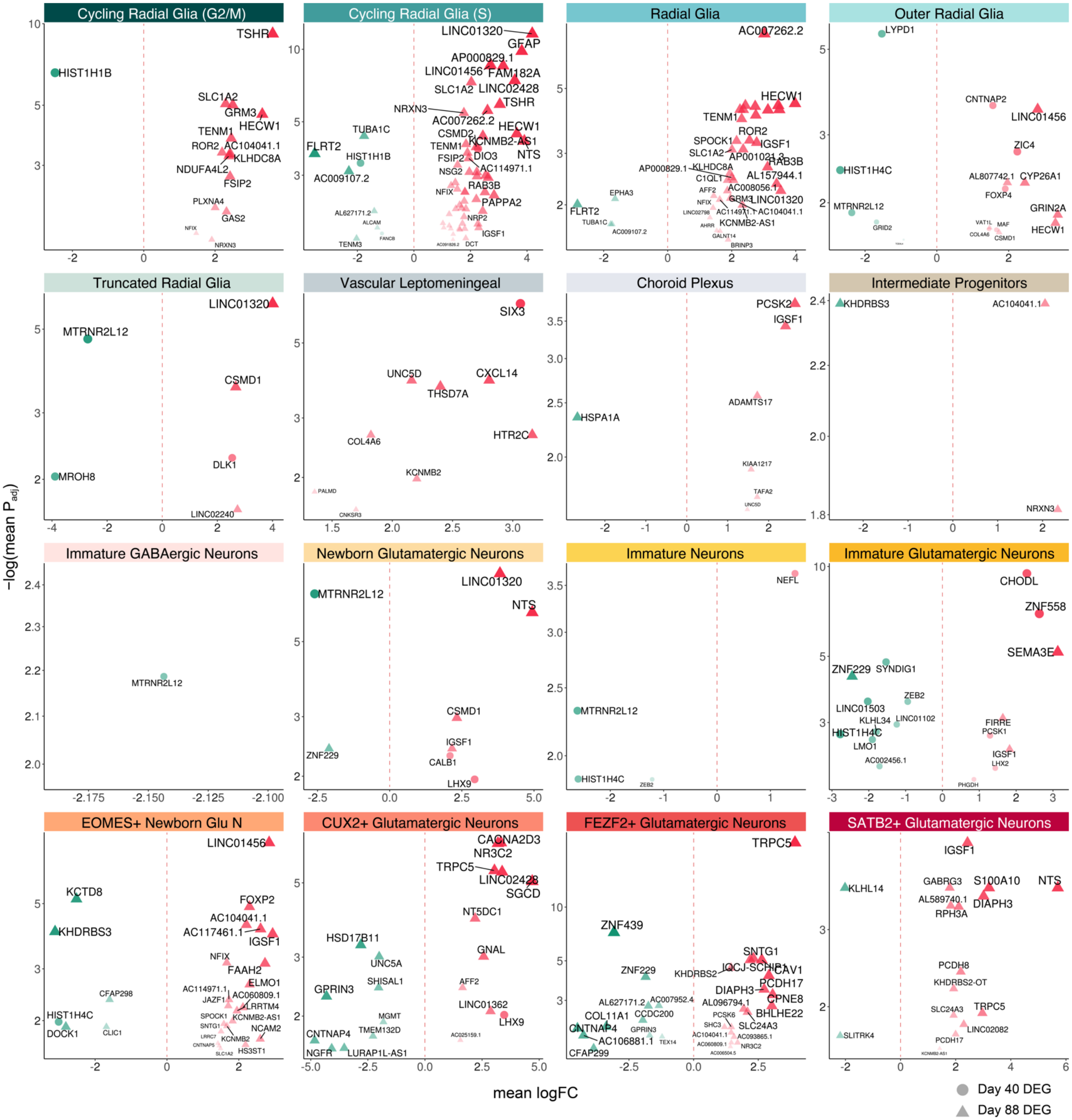


**Figure S9. All differentially expressed genes.** A volcano plot of the 277 significant differentially expressed genes (DEGs) by cell type. The x axis denotes mean logFC of the DEG between standard Monocle3 and pseudobulk DESeq2 methods. The y axis denotes the mean adjusted *p* value of the DEG between methods. y axis is shown on a $\log_{10}$ scale with a cutoff of $p_{adj} < 0.05$ for both DEG identification methods used. Size and alpha correlates to a combination of lowest mean $p_{adj}$ and highest mean absolute logFC. Shape denotes the dataset time point at which the DEG was identified. Red color indicates the gene is upregulated and green downregulated in PACS1$^{(+/R203W)}$ cells.

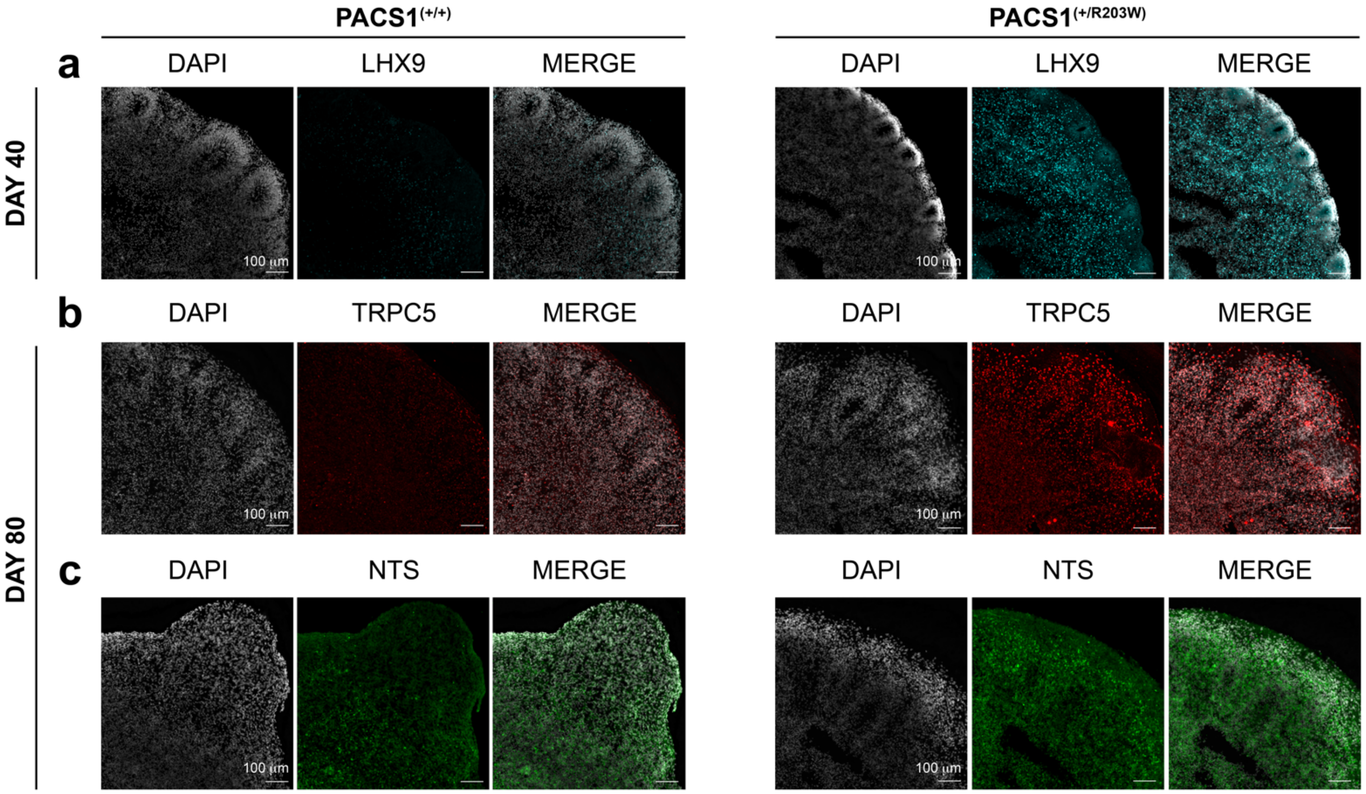


**Figure S10: Protein expression of top upregulated DEGs in PACS1$^{(+/R203W)}$ organoids identified with single-cell RNA sequencing. a)** Expression of LHX9 in a CRISPR A2 day 40 PACS1$^{(+/+)}$ organoid and an A1 PACS1$^{(+/R203W)}$ organoid. Scale bar = 100 µm. Single-cell RNA sequencing (scRNAseq) analysis predicted upregulation of LHX9 in PACS1$^{(+/R203W)}$ day 40 organoids in bulk, newborn glutamatergic neurons, and CUX2+ glutamatergic neurons. **b)** Expression of TRPC5 in a CRISPR A1 day 80 PACS1$^{(+/+)}$ organoid and an A1 PACS1$^{(+/R203W)}$ organoid. Scale bar = 100 µm. scRNAseq analysis predicted upregulation of *TRPC5* in PACS1$^{(+/R203W)}$ day 88 organoids in bulk and all three mature glutamatergic neuron groups. **c)** Expression of NTS in a C2 day 80 PACS1$^{(+/+)}$ organoid and an A1 PACS1$^{(+/R203W)}$ organoid. Scale bar = 100 µm. scRNAseq analysis predicted upregulation of *NTS* in PACS1$^{(+/R203W)}$ day 88 in bulk, newborn glutamatergic neurons, cycling radial glia (S), and SATB2+ glutamatergic neurons.

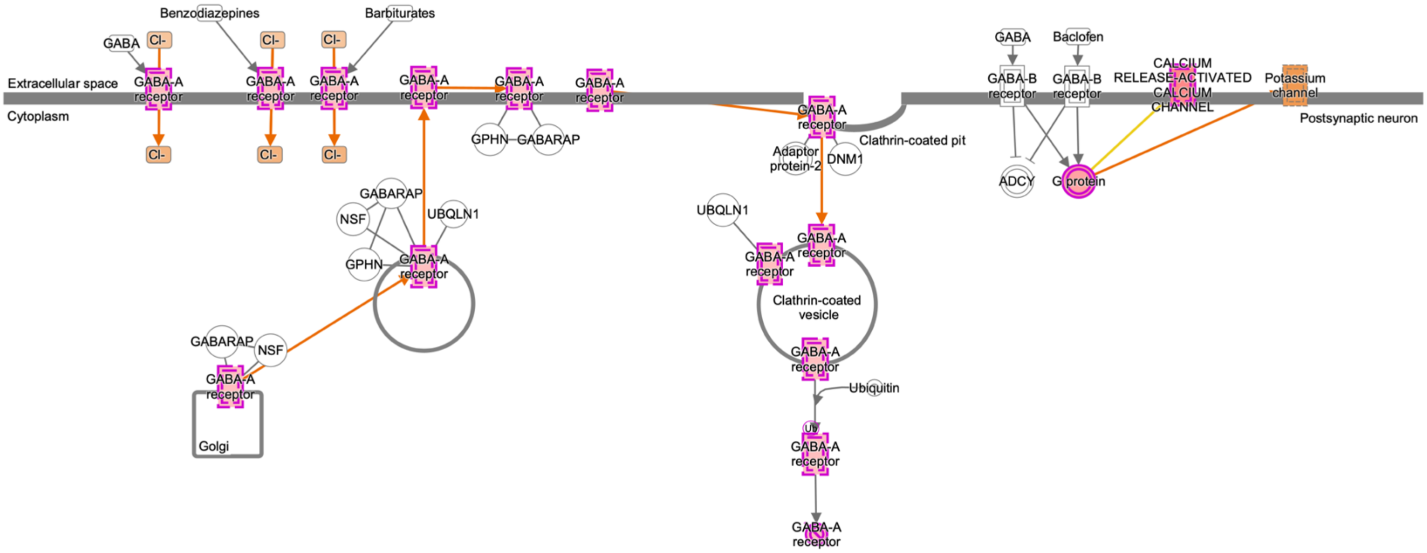


**Figure S11. Ingenuity Pathway Analysis predicts differentially expressed genes in PACS1$^{(+/R203W)}$ neurons result in increased GABA-A receptor activity. a)** Pathway diagram generated by Qiagen Ingenuity Pathway analysis predicting upregulation of GABA-A receptor activity in PACS1$^{(+/R203W)}$ neurons ($p_{adj}$ = 0.016).

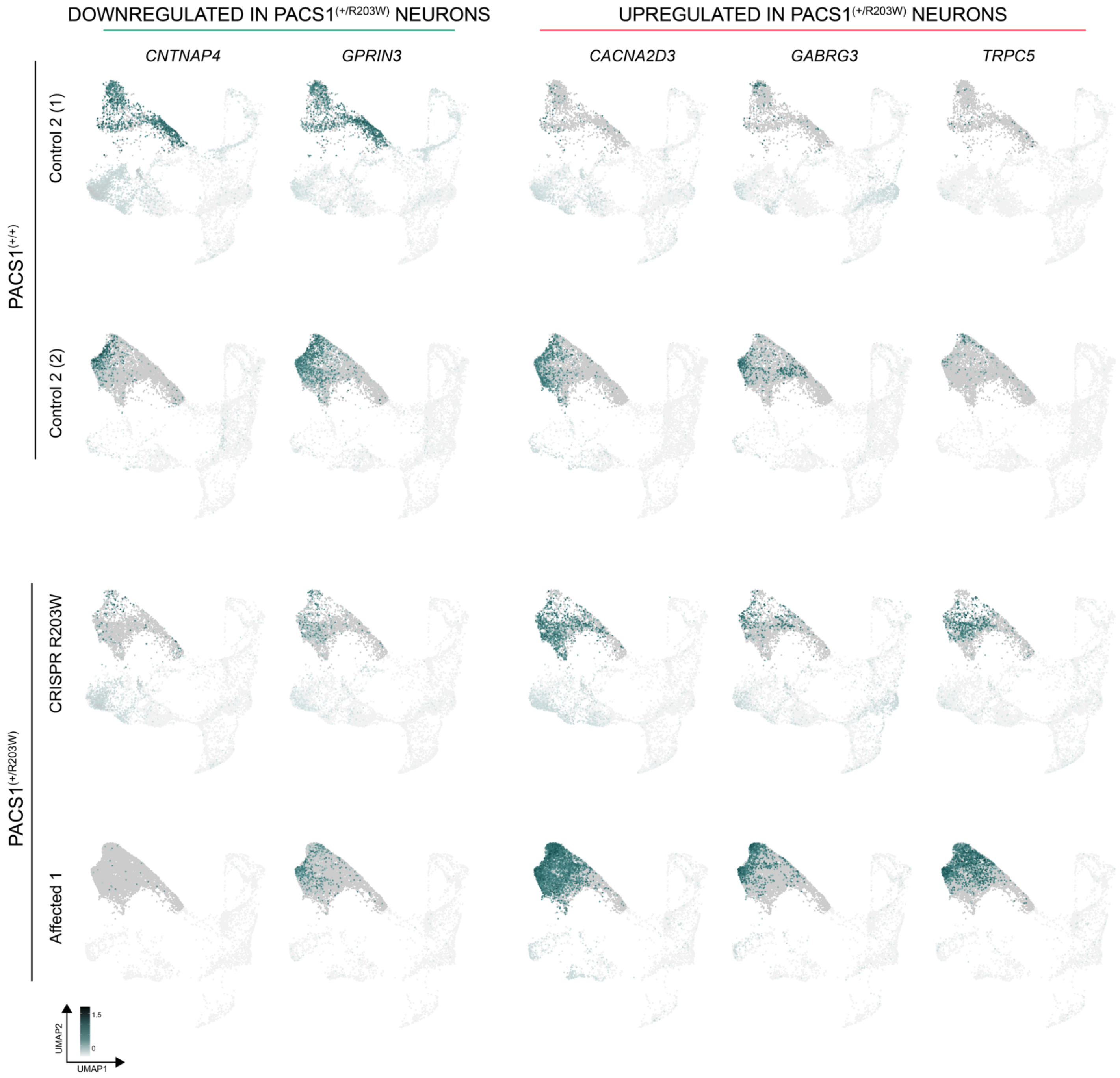


**Figure S12. Top differentially expressed genes by sample.** Feature plots of top differentially expressed genes (DEGs) in PACS1$^{(+/R203W)}$ mature neurons implicated in Ingenuity Pathway analysis. Expression is shown across each day 88 organoid sequenced. Cells have been down-sampled so the same number is shown for each genotype. Scale is in units of $\log_{10}$(value + 0.1). The opacity of non-mature neuron groups has been decreased in order to highlight the cell types of interest. Data is shown from two C2, one CRISPR R203W, and one A1 day 88 organoid from one independent differentiation).

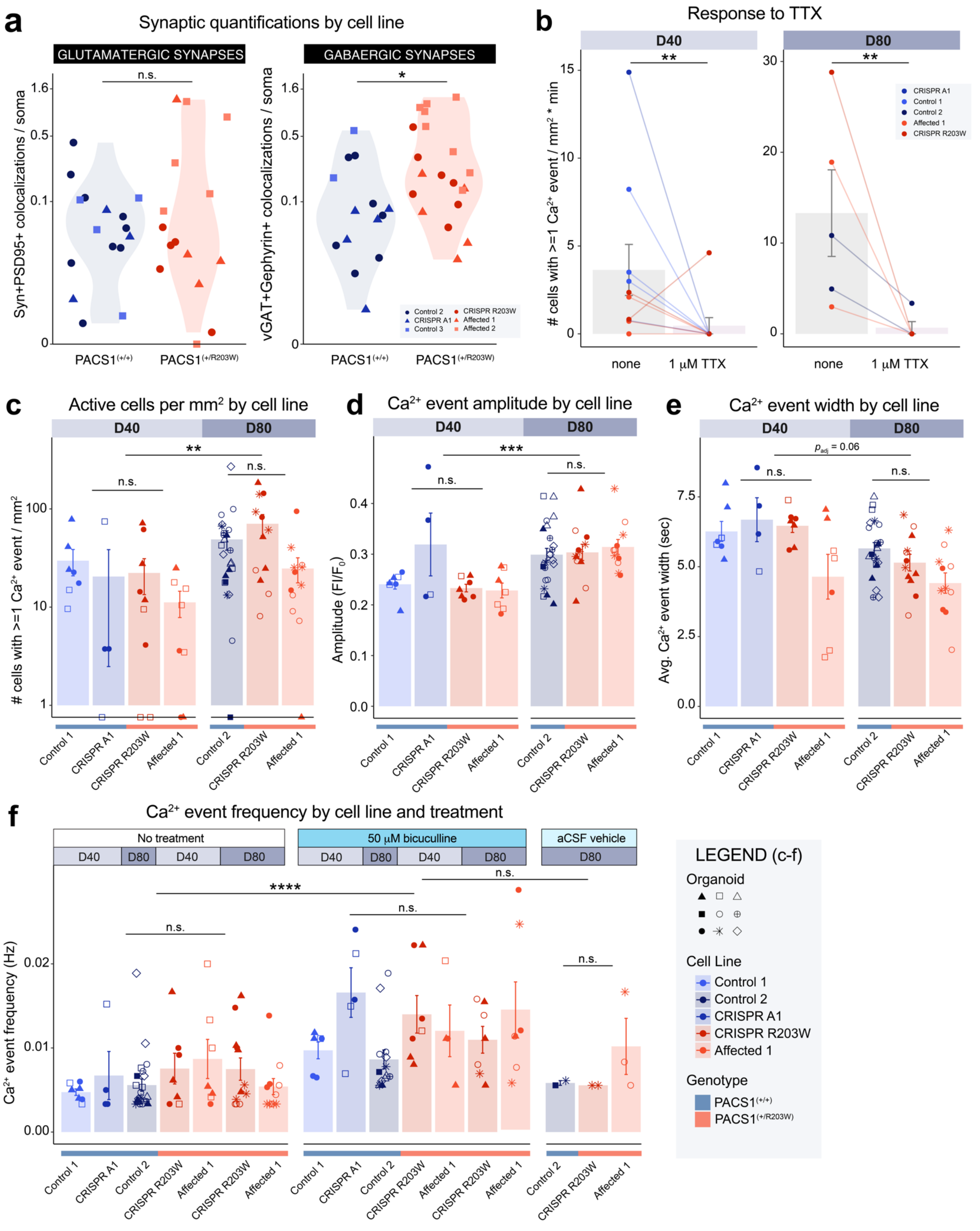
a
Synaptic quantifications by cell line
GLUTAMATERGIC SYNAPSES
GABAERGIC SYNAPSES
Syn+PSD95+ colocalizations / soma
vGAT+Gephyrin+ colocalizations / soma
n.s.
*
PACS1(+/+)
PACS1(+/R203W)
Control 2
CRISPR A1
Control 3
CRISPR R203W
Affected 1
Affected 2
b
Response to TTX
D40
D80
### cells with >=1 Ca2+ event / mm2 * min
none
1 μM TTX
CRISPR A1
Control 1
Control 2
Affected 1
CRISPR R203W
c
Active cells per mm2 by cell line
### cells with >=1 Ca2+ event / mm2
d
Ca2+ event amplitude by cell line
Amplitude (FI/F0)
e
Ca2+ event width by cell line
Avg. Ca2+ event width (sec)
padj = 0.06
Control 1
CRISPR A1
CRISPR R203W
Affected 1
Control 2
f
Ca2+ event frequency by cell line and treatment
No treatment
50 μM bicuculline
aCSF vehicle
Ca2+ event frequency (Hz)
****
LEGEND (c-f)
Organoid
Cell Line
Control 1
Control 2
CRISPR A1
CRISPR R203W
Affected 1
Genotype
PACS1(+/+)
PACS1(+/R203W)

**Figure S13. PACS1$^{(+/R203W)}$ organoids have increased GABAergic synaptic density but no differences in neural activity detected by calcium imaging. a)** Data from **Fig. 4b-c** shown by cell line. In the left panel, dots represent number of SYN1$^{+}$PSD95$^{+}$ colocalizations per soma in day 80 organoid sections. y axis is on a $\log_2$ scale. Data is from $n$ = 16 total sections across two C2, one C3, and one CRISPR A1 PACS1$^{(+/+)}$ organoids and $n$ = 15 total sections across two CRISPR R203W, one A1, and two A2 PACS1$^{(+/R203W)}$ organoids from three independent differentiations. See **Table S1**. $U$=118, $p$ = 0.953 using a two-sided Mann-Whitney test. In the right panel, dots are number of vGAT$^{+}$Gephyrin$^{+}$ colocalizations in day 80 organoid sections shown in **Fig. 4c** by cell line. y axis is on a $\log_2$ scale. Data is from $n$ = 14 sections across two C2, one C3, and one CRISPR A1 PACS1$^{(+/+)}$ organoids and $n$ = 20 sections across two CRISPR R203W, one A1, and two A2 PACS1$^{(+/R203W)}$ organoids from two independent differentiations. See **Table S1**. $U$=71, $p$ = 0.015 calculated using a two-sided Mann-Whitney test. **b)** Number of cells with at least one $Ca^{2+}$ event per mm$^2$ of an organoid section after exposure to 1 µM tetrodotoxin (TTX) per minute of recording. TTX has an effect on number of active cells ($W$ = 84, $p$ = 0.008) using a Wilcoxon signed rank test on the difference between paired values ($p_{\text{Shapiro}}$ = 0.012). Data is from $n$ = 5 PACS1$^{(+/+)}$ D40 organoids (three C1, two CRISPR A1), $n$ = 5 PACS1$^{(+/R203W)}$ D40 organoids (three A1, two CRISPR R203W), $n$ = 2 C2 PACS1$^{(+/+)}$ D80 organoids, and $n$ = 2 PACS1$^{(+/R203W)}$ D80 organoids (one A1, one CRISPR R203W). **c)** Results from **Fig. 4e** shown by cell line. Bar chart showing mean ± SEM of number of active cells per mm$^2$ of an organoid section. Scale is $\log_{10}$ for clarity. There is an effect of day ($F_{1,23}$=8.576, $p_{\text{adj}}$ = 0.008) but not genotype ($F_{1,23}$=0.482, $p_{\text{adj}}$ = 0.495) using a two-way ANOVA test with a Tukey HSD correction on log-transformed values. Shapes represent values recorded from distinct $n$ = 5 Day 40 PACS1$^{(+/+)}$ organoids (three C1 and two CRISPR A1), $n$ = 6 Day 40 PACS1$^{(+/R203W)}$ organoids (three CRISPR R203W and three A1), $n$ = 8 Day C2 80 PACS1$^{(+/+)}$ organoids, and $n$ = 7 Day 80 PACS1$^{(+/R203W)}$ organoids (four CRISPR R203W and three A1) from two independent differentiations for panels **c-e**. **d)** Data from **Fig. 4f** shown by cell line. Bar chart displaying the mean ± SEM of $Ca^{2+}$ event amplitude. There is an effect of day ($F_{1,23}$=15.481, $p_{\text{adj}}$=6.815E-04) but not genotype ($F_{1,23}$=0.484, $p_{\text{adj}}$=0.494) using a two-way ANOVA test with a Tukey HSD correction. **e)** Data from **Fig. 4g** shown by cell line. Bar chart displaying the mean ± SEM of $Ca^{2+}$ event width. There may be an effect of day ($F_{1,23}$=3.811, $p_{\text{adj}}$=0.064) but not genotype ($F_{1,23}$=2.648, $p_{\text{adj}}$=0.117) using a two-way ANOVA test with a Tukey HSD correction. **f)** Bar chart displaying mean ± SEM of average frequency of $Ca^{2+}$ events for each cell line in **Fig. 4h**, as well as any non-paired values. There is an effect of 50 µM bicuculline (BCU) exposure on $Ca^{2+}$ event frequency ($H_1$= 18.468, $p_{\text{adj}}$ = 8.6E-05) but no other parameters tested using one-sided Kruskal Wallis tests followed by Benjamini Hochberg correction ($p_{\text{Shapiro}}$ = 6.6E-04). Shapes represent values recorded from distinct $n$ = 5 Day 40 PACS1$^{(+/+)}$ organoids (three C1 and two CRISPR A1), $n$ = 6 Day 40 PACS1$^{(+/R203W)}$ organoids (three CRISPR R203W and three A1), $n$ = 8 Day C2 80 PACS1$^{(+/+)}$ organoids, and $n$ = 8 Day 80

PACS1$^{(+/R203W)}$ organoids (four CRISPR R203W and four A1) from two independent differentiations. Legend also applies to panels c-e.

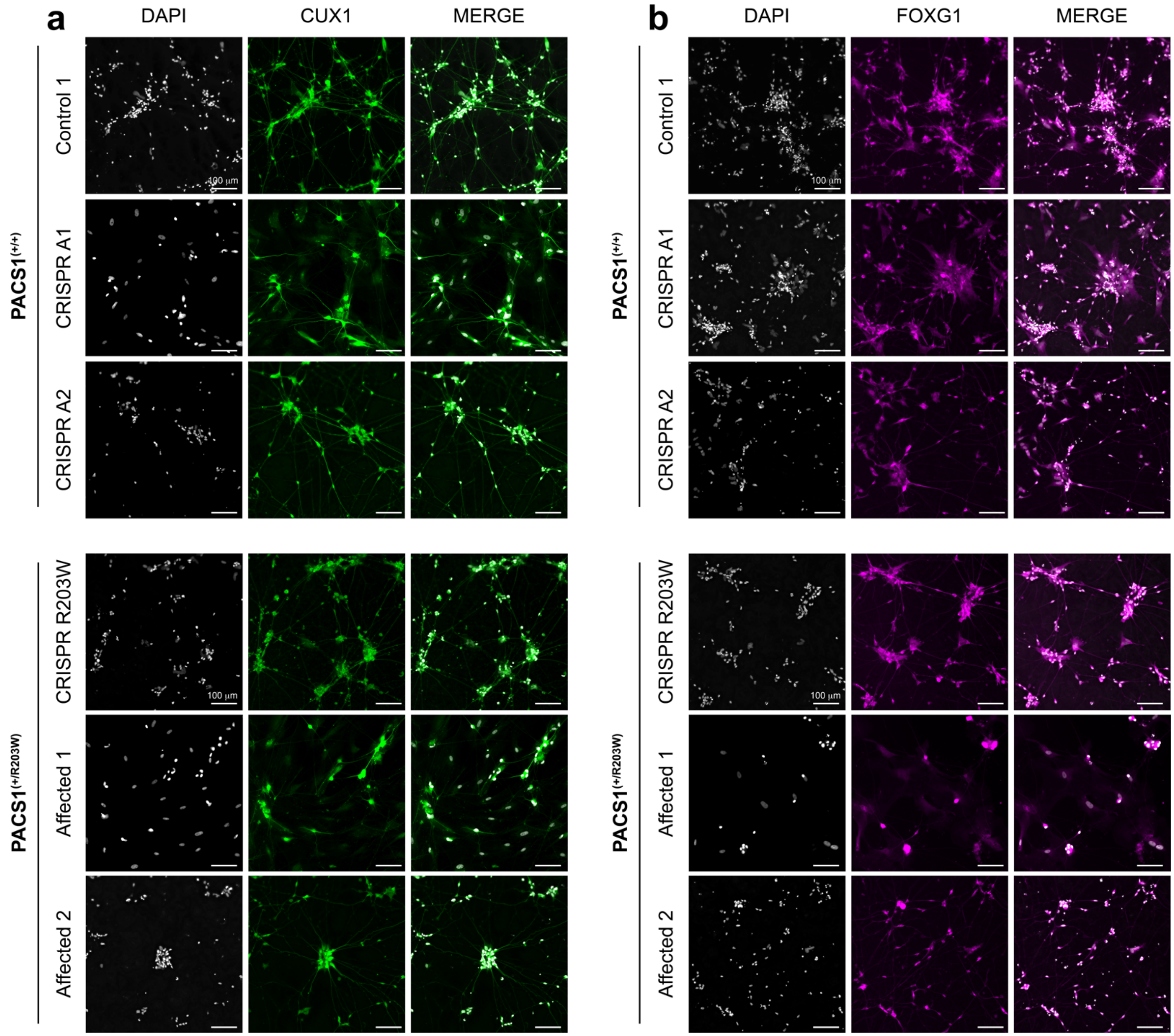


**Figure S14. Neurons generated from iPSCs by over-expressing NEUROG2 express forebrain markers. a)** Both PACS1$^{(+/+)}$ and PACS1$^{(+/R203W)}$ ~D50 NEUROG2 neurons show expression of layer II/III neuron marker CUX1. Scale bar = 100 µm. **b)** Both PACS1$^{(+/+)}$ and PACS1$^{(+/R203W)}$ ~D50 NEUROG2 neurons show expression of forebrain neuronal transcription factor FOXG1. Scale bar = 100 µm.

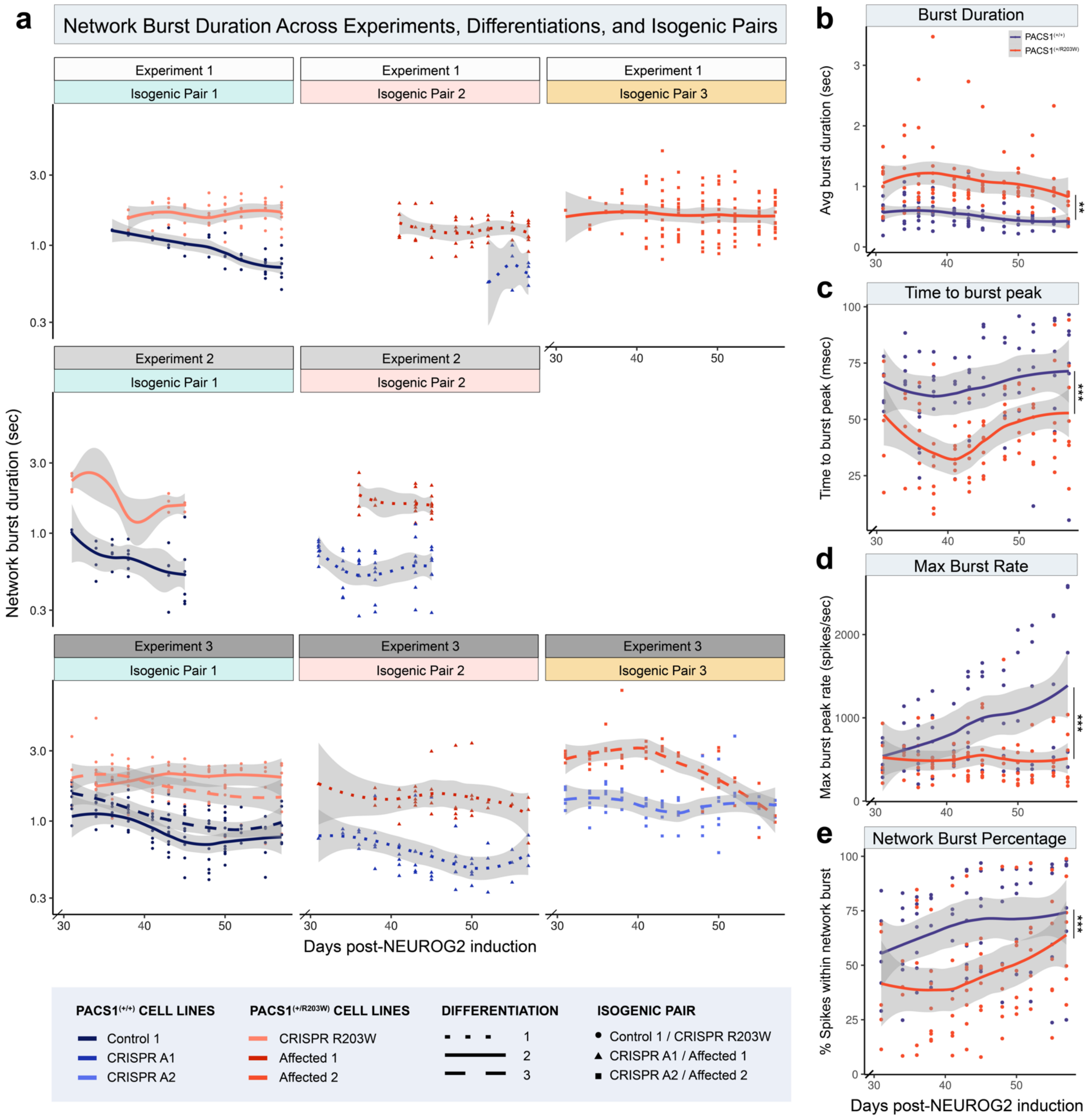


**Figure S15. PACS1$^{(+/R203W)}$ neurons consistently display a prolonged network burst duration. a)** Dots show multielectrode array (MEA) well averages of network burst duration for each experiment, differentiation, and isogenic pair. The x axis has been abbreviated to days 30-57 as network bursts do not occur before day 30. Dots represent biological replicate averages from MEA wells. The trend line indicates genotype averages with a 95% confidence interval. $n$ = 7 PACS1$^{(+/+)}$ lines (three C1, three CRISPR A1, and one CRISPR A2) and $n$ = 7 PACS1$^{(+/R203W)}$ lines (three CRISPR R203W, three A1, and one A2) from three independent experiments. See **Table S1**. Missing values are either due to either: the neurons not yet being mature enough to elicit network bursting activity; the line not being included in the MEA experiment; or omission of the data upon observation of excessive clumping or other indicators of poor viability. **b-e)** Further MEA metrics for neurons days 30-57 of differentiation. **b)** Average burst duration in seconds. Genotype had a significant effect on burst duration ($F_{0,2} = 5.261$, $p = 0.006$). **c)** Time from the start to the peak of the average network burst. Genotype had a significant effect on time to burst peak ($F_{0,2} = 26.686$, $p = 8.48E\text{-}11$). **d)** Maximum number of spikes per second in the average network burst. Genotype had a significant effect on max burst rate ($F_{0,2} = 27.413$, $p = 4.89E\text{-}11$). **e)** Percentage of total spikes that occur within network bursts. Genotype had a significant effect on network burst percentage ($F_{0,2} = 14.705$, $p = 1.30E\text{-}06$). Individual time points are significant at days 38 – 43 of differentiation.

#### SUPPLEMENTARY TABLES

| | PACS1(+/+) | | | | | PACS1(+/R203W) | | | | | |
|---|---|---|---|---|---|---|---|---|---|---|---|
| Cell line | Control 1 | Control 2 | Control 3 | CRISPR A1 | CRISPR A2 | CRISPR R203W | Affected 1 | Affected 2 | | | |
| Identifaction/Line of Origin | GM03651 | GM03652 | GM27160 | GM27161 | GM27159 | GM03651 | GM27161 | GM27159 | ***n* represents** | **Independent Differentiations** | **Applicable Figures** |
| Sex | F | M | F | M | F | F | M | F | | | |
| NPC BRDU assay | 1 | 1 | 1 | 0 | 0 | 1 | 1 | 1 | Cell line averages / experiment | 2 | Fig S5c-d |
| NPC TUNEL assay | 4 | 0 | 2 | 0 | 0 | 6 | 2 | 4 | Cell line averages / experiment | 2 | Fig S5e-f |
| FOXG1/TBR1/SATB2 D40 org | 3 | 0 | 0 | 0 | 0 | 1 | 2 | 0 | One organoid | 4+ | Fig 1d, S4d |
| FOXG1/TBR1/SATB2 D80 org | 0 | 2 | 0 | 1 | 0 | 2 | 1 | 0 | One organoid | 3+ | Fig 1d, S4d |
| CTIP2/SOX2 D40 org | 5 | 0 | 1 | 1 | 1 | 1 | 5 | 1 | One organoid | 4+ | Fig 1e, S4e |
| CTIP2/SOX2 D80 org | 2 | 3 | 1 | 1 | 0 | 3 | 4 | 0 | One organoid | 3+ | Fig 1e, S4e |
| PACS1 western blot D40 org | 2 | 3 | 0 | 2 | 0 | 4 | 4 | 0 | One organoid | 1 | Fig 1f-g |
| scRNAseq day 40 org | 1 | 0 | 1 | 0 | 0 | 1 | 1 | 1* | One organoid | 3 | Figs 2-3, S6-S9 |
| scRNAseq day 88 org | 0 | 2 | 0 | 0 | 0 | 1 | 1 | 0 | One organoid | 1 | Figs 2-3, S6-S9 |
| Syn1/PSD95 D79 org | 0 | 2 | 1 | 1 | 0 | 2 | 1 | 2 | One organoid section | 3 | Fig 4b, S13a |
| vGAT/GPHN D79 org | 0 | 2 | 1 | 1 | 0 | 2 | 1 | 2 | One organoid section | 2+ | Fig 4c, S13a |
| Calcium imaging D40 org | 3 | 0 | 0 | 2 | 0 | 3 | 3 | 0 | One organoid | 1 | Fig 4d-i, S13b-f |
| Calcium imaging D80 org | 0 | 8 | 0 | 0 | 0 | 4 | 4 | 0 | One organoid | 1 | Fig 4d-i, S13b-f |
| Sholl analysis NGN2 neurons | 1 | 0 | 0 | 2 | 1 | 1 | 2 | 1 | Cell line averages / experiment | 2 | Fig 5d-g |
| MEAs NGN2 neurons | 3 | 0 | 0 | 3 | 1 | 3 | 3 | 2 | Cell line averages / experiment | 3 | Fig 5h-o, S15 |

**Table S1. Number of samples from each cell line contributing to each experiment.**

Distribution of samples from each cell line used for each assay. * Indicates a specific instance in which two technical replicates that returned a low number of sequenced cells were combined and treated as one organoid. + indicates at least that many differentiations were used in quantifications. Please note that independent differentiations indicates the total number of differentiations present across all samples in a given experiment, but does not necessarily indicate the experiment was repeated that many number of times with the exact same lines.

**Table S2. The complete list of differentially expressed genes.** (Separate file)

The complete list of differentially expressed genes (DEGs) between PACS1$^{(+/+)}$ and PACS1$^{(+/R203W)}$ organoids.
